# Solid Ionic Matrices applied via Low-Temperature Evaporation enable High-Resolution and Sensitive MALDI Imaging of Metabolites

**DOI:** 10.1101/2025.06.23.661081

**Authors:** Toufik Mahamdi, Nadia Moltó, Mireia Arasa, Roger Giné, Jordi Rofes, María García-Altares, Maria Vinaixa, Oscar Yanes

## Abstract

Matrix-Assisted Laser Desorption/Ionization Mass Spectrometry Imaging (MALDI-MSI) has become a key technology for spatially resolved molecular analysis. However, imaging of low molecular weight compounds remains challenging due to spectral interferences in the low *m/z* range caused by conventional organic matrices. Ionic matrices (IMs), in contrast, reduce matrix-related background signals, improve spectral reproducibility, and enhance sensitivity for detecting metabolites and lipids, but their use in MALDI-MSI has so far been limited, primarily focusing on lipid imaging above *m/z* 500. Here, we present a novel approach combining solid ionic matrices (ISMs) with low-temperature thermal evaporation (LTE) to enhance MALDI-MSI performance for small metabolites. ISMs based on α-cyano-4-hydroxycinnamic acid (CHCA) and 2,5-dihydroxybenzoic acid (DHB) provided superior sensitivity, with CHCA + N,N-diethylaniline enabling the detection and annotation of 73 small metabolites (*m/z* <500) in mouse brain at 20 µm spatial resolution. When applied to the pancreas, this matrix enabled the first specific localization of melatonin and 3-iodotyrosine within the islets of Langerhans, opening new avenues for spatial metabolomics in endocrine research.

## 1. Introduction

Spatial metabolomics, enabled by Matrix-Assisted Laser Desorption/Ionization Mass Spectrometry Imaging (MALDI-MSI), has become a transformative technology for detecting and mapping the spatial distribution of metabolites, lipids, and other small molecules within biological tissues [1]. By preserving the spatial context of molecular information—often at cellular or even subcellular resolution [2]—MALDI-MSI offers unique insights into tissue heterogeneity, biochemical processes, and disease mechanisms, playing a key role in biomarker discovery and spatially resolved molecular phenotyping [3].

A key component of MALDI-MSI is the matrix compound, which is co-crystallized with the analyte on the tissue surface. The matrix absorbs laser energy and mediates the desorption and ionization of analyte molecules, playing a critical role in determining ionization efficiency, sensitivity, reproducibility, and the overall range of detectable metabolites [4–8]. However, conventional organic matrices—such as those based on cinnamic acids (e.g., α-cyano-4-hydroxycinnamic acid, CHCA) and benzoic acids (e.g., 2,5-dihydroxybenzoic acid, DHB)— generate substantial background signals in the low-mass range, typically below *m/z* 500. These matrix-derived ions can interfere with the ionization, detection, and accurate quantification of small molecules [9].

To address these limitations and improve MALDI-MSI performance for metabolite analysis, a range of strategies have been developed. These include the incorporation of matrix additives [9], the design of novel matrices with tailored physicochemical properties [10–13], the use of high molecular weight matrices to reduce background interference, and on-tissue derivatization protocols that chemically shift analytes to higher m/z values [14–16]. Additional approaches involve the use of metal nanoparticles [17] and optimized matrix deposition techniques [18–20], all aimed at enhancing ionization efficiency and signal-to-noise ratio for small molecules detection.

In this context, ionic matrices (IMs) [21] have also emerged as a promising alternative to conventional organic matrices. IMs—available in either liquid (ionic liquid matrices, ILMs) or solid (ionic solid matrices, ISMs) forms—are organic salts formed through acid–base reactions, typically involving equimolar mixtures of standard acidic MALDI matrices (e.g., DHB or CHCA) and organic bases such as amines [22,23]. IMs offer several advantages over conventional MALDI matrices. They exhibit greater thermal and vacuum stability and significantly reduce matrix-related background signals. These properties lead to cleaner mass spectra, improved reproducibility, and enhanced sensitivity for detecting small molecules, including both metabolites and lipids [24].

Despite these benefits, the application of IMs in MALDI-MSI remains limited [25]. So far, studies have primarily focused on liquid IMs, applied via wet deposition techniques such as spraying, and have largely been restricted to lipid imaging in the *m/z* >500 range [26–29].

To enhance small molecule detection by MALDI-MSI, we combined ionic solid matrices (ISMs) with low-temperature thermal evaporation (LTE), a dry matrix deposition method that improves ionization efficiency, minimizes analyte diffusion, and enhances spatial resolution. We evaluated three ISMs—CHCA+DEA, CHCA+DMA, and DHB+ANI—against conventional matrices CHCA and DHB in MALDI-MSI analyses of mouse brain and liver tissues. All ISMs demonstrated superior performance, enabling enhanced detection of low molecular weight metabolites while substantially reducing matrix-related background signals. Notably, CHCA+DEA exhibited the most pronounced improvements, supporting high-resolution imaging of both brain and pancreatic tissues. This matrix facilitated the annotation of 73 endogenous metabolites in mouse brain sections, including several that delineated pancreatic islets with high contrast and specificity.

## 2. Materials and Methods

### 2.1. Reagent and Chemicals

Ethanol (HPLC grade) was obtained from Scharlab S.L. The following compounds were purchased from Sigma-Aldrich: α-cyano-4-hydroxycinnamic acid (CHCA), 2,5-dihydroxybenzoic acid (DHB), aniline (ANI), N,N-diethylaniline (DEA), and N,N-dimethylaniline (DMA). Analytical standards of alanine, creatine, cholesterol, glutamic acid, glutamine, glutathione, 4-hydroxybenzoic acid (4-HB), 3-indoleacetic acid (3-IAA), 3-iodotyrosine, melatonin, 3-methylglutaryl carnitine, kynurenic acid (KYNA), taurine, and tyrosine were also purchased from Sigma-Aldrich. Microscope slides were obtained from Epredia, and conductive indium tin oxide (ITO)-coated slides were sourced from Bruker.

### 2.2. Tissue Sampling

Brain, liver, and pancreas tissues were harvested from healthy mice, flash-frozen on dry ice, and stored at –80°C until further processing. Tissue sections were prepared using a CryoStar NX50 cryostat (Thermo Fisher Scientific) at –20°C, sectioned at a thickness of 10 µm. Sections were mounted onto ITO-coated microscope slides and immediately stored at –80°C until analysis.

### 2.3. Synthesis of Solid Ionic Matrices

For the preparation of ionic matrices (IM), a stock solution of the organic matrix was first prepared by dissolving 1 g of either DHB or CHCA in 50 mL of ethanol. An equimolar amount of the base—aniline (ANI), N,N-diethylaniline (DEA), or N,N-dimethylaniline (DMA)—was then added to the matrix solution to form the corresponding IM solution. The mixture was vortexed for 15 minutes to ensure homogeneity.

To obtain solid IMs, the solvent was removed by subjecting the solutions to a nitrogen gas stream using a solid-phase extraction apparatus. To expedite solvent evaporation, each IM solution was divided into four 50 mL glass bottles. After complete solvent removal, the resulting dried IMs were collected into Eppendorf tubes for subsequent LTE deposition.

### 2.4. Matrix Application

Matrix coating was performed using a NanoPVD-T15A thermal evaporator (Moorfield Nanotechnology, United Kingdom), following the low-temperature thermal evaporation (LTE) protocol described by Mahamdi et al. [30]. A total of 300 mg of matrix material—either conventional or solid ionic matrices (IMs)—was placed into the crucible of the LTE source, while the tissue section was mounted on the substrate holder.

After evacuating the chamber to a base pressure of 5 × 10⁻⁵ mbar, the initial temperature was set to 60 °C. The system was allowed to stabilize, defined as the point at which the power supply unit dropped to zero. Subsequently, the temperature was gradually increased in 5 °C increments until the target evaporation temperature for the specific matrix was reached (Supplementary Table S1). Once at this temperature, the substrate shutter was opened, allowing the vaporized matrix to coat the tissue surface. Deposition thickness was precisely controlled and monitored throughout the process, reaching 15,000 Å for DHB and DHB-based solid IMs, and 9,000 Å for CHCA and CHCA-based solid IMs.

### 2.5. H&E Staining of Tissue Sections

A standard hematoxylin and eosin (H&E) staining protocol was applied to consecutive sections adjacent to those analyzed by MALDI-MSI. The staining procedure was as follows: tissue sections were stained with hematoxylin for 30 seconds, followed by sequential washes with tap water, 1% acetic acid, and distilled water for 10 seconds each. The sections were then stained with eosin for 20 seconds and dehydrated through graded ethanol solutions (70%, 96%, and 100%) for 10 seconds each. Finally, the tissues were cleared in Neo-Clear for 40 seconds.

The stained sections were mounted with coverslips using Cytoseal XYL (Richard-Allan Scientific, Kalamazoo, MI, USA) and allowed to dry at room temperature. Histological images were acquired at 40x magnification using the Ocus®40 microscope slide scanner (Grundium, Finland).

### 2.6. MALDI-MSI and MALDI-MS/MS acquisition

MALDI-MSI and MS/MS analyses were performed using a dual-ion funnel MALDI/ESI injector (Spectroglyph, Kennewick, WA) coupled to an Orbitrap Exploris 120 mass spectrometer (Thermo Fisher Scientific). For MSI acquisitions, spatial step sizes of 20 µm and 30 µm were used. Spectra were acquired over the *m/z* range 50–600 Da at a resolving power of 60,000 (FWHM at *m/z* 200).

MSI datasets were processed, and ion images reconstructed and visualized using the rMSI2 R package (https://github.com/prafols/rMSI2) [31]. Ion annotations were performed using rMSIannotation [32] and tentatively identified by matching against the Human Metabolome Database (HMDB) (https://www.hmdb.ca/), considering the following adducts: [M+H]^+^, [M−H₂O+H]^+^, [M+Na]^+^, [M+K]^+^, [M+2Na−H]^+^, and [M+2K−H]^+^, with a mass tolerance of 10 ppm.

For MS/MS experiments, collision-induced dissociation (CID) was performed with a normalized collision energy of 30%.

## 3. Results and Discussion

### 3.1. Evaluation of ionic solid matrices (ISM) for MALDI-MSI

Six ionic matrices (IMs) were synthesized by combining two conventional acidic MALDI matrices—CHCA and DHB—with three different organic bases: aniline (ANI), N,N-diethylaniline (DEA), and N,N-dimethylaniline (DMA). This resulted in six IM formulations: CHCA+ANI, CHCA+DEA, CHCA+DMA, DHB+ANI, DHB+DEA, and DHB+DMA. However, DHB+DEA and DHB+DMA did not yield solid products upon solvent evaporation and were therefore excluded from further analysis (Supplementary Table S1).

We next evaluated the suitability of the solid IMs (ISMs) for deposition via low-temperature thermal evaporation (LTE). The evaporation temperature for each ISM was optimized individually (Supplementary Table S1). Compared to their conventional counterparts—CHCA (100 °C) and DHB (80 °C)—the solid IMs required evaporation temperatures approximately 10–15 °C higher. This increase is likely due to the incorporation of base components in CHCA and DHB, which have lower volatility and require more heat for effective evaporation.

Given that the acidic and basic components in ISMs have different volatility profiles, we assessed whether the optimized LTE conditions were sufficient to co-deposit both components onto mouse brain and liver tissue surfaces. To do so, we performed MALDI-MSI analyses of the matrix coatings, confirming the presence of the protonated and sodium-adduct ions of the base molecules (ANI, DEA, DMA) of all ISM formulations (Supplementary Fig. S1). These results confirm that both components were successfully deposited and that the resulting matrix coatings retained the characteristic properties of ionic matrices.

Next, we assessed the performance of ISMs deposited via LTE for the ionization and detection of small metabolites within the *m/z* 50–600 Da range. We began by analyzing the total ion count (TIC) per pixel across all tissue samples. In both brain and liver tissues, ISMs consistently outperformed conventional matrices, producing higher TIC values. Among the CHCA-based ISMs, CHCA+DEA generated the highest TIC in brain tissue (Figure 1a), while CHCA+DMA yielded the strongest signal in liver samples (Figure 1b).

**Figure 1:**
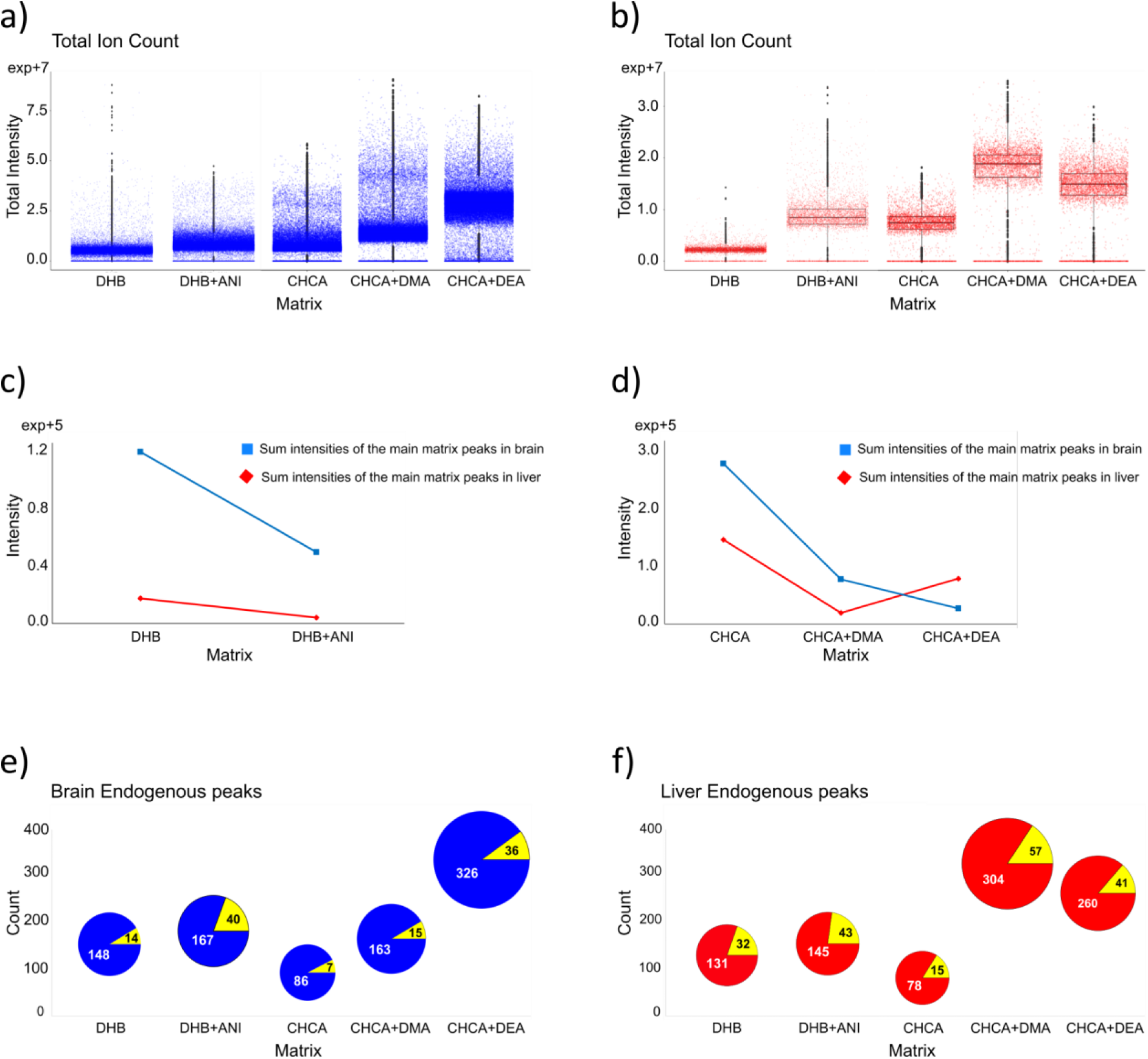
Comparative MALDI-MSI performance of DHB, DHB+ANI, CHCA, CHCA+DMA, and CHCA+DEA matrices. (a, b) Boxplots of total ion count (TIC) intensities from mouse brain (a) and liver (b) tissue. (c, d) Sum intensities of key matrix-related ions for DHB-based (c) and CHCA-based (d) matrices detected in brain (blue) and liver (red) tissues. (e, f) Pie charts showing the proportion of tissue-related peaks annotated using HMDB (blue or red: annotated ions; yellow: no hit) in brain (e) and liver (f).

To investigate the source of these increased TIC values, we first examined the formation of the predominant matrix-related ion clusters associated with each ISM. For DHB and DHB-based ISMs, key signals included *m/z* 137.0239 ([M–H₂O+H]⁺), *m/z* 155.0345 ([M+H]⁺), *m/z* 273.0399 ([2M–2H₂O+H]⁺), and *m/z* 409.0559 ([3M–3H₂O+H]⁺) (Supplementary Fig. S2). For CHCA and CHCA-based ISMs, the main matrix-related ions were *m/z* 146.0613 ([M–CO+H]⁺), *m/z* 172.0394 ([M–H₂O+H]⁺), *m/z* 190.0511 ([M+H]⁺), *m/z* 212.0340 ([M+Na]⁺), and *m/z* 228.0067 ([M+K]⁺) (Supplementary Fig. S3). In all cases, regardless of tissue type, the signal intensity of matrix-related ions was markedly reduced in samples prepared with ISMs, resulting in lower overall matrix background (Figure 1c and 1d). These results strongly suggested that the enhanced TIC observed with ISMs was primarily due to improved ionization of endogenous metabolites rather than contributions from matrix-related ions. To confirm this, we quantified the number of biologically relevant signals detected in each condition. By comparing spectra acquired from tissue regions with those from adjacent matrix-only areas, we filtered out matrix-derived peaks and retained only tissue-specific ions. These were then annotated using the rMSI2 R package and matched against the Human Metabolome Database (HMDB) using a defined set of matrix-specific adducts (see Methods for details).

Using the conventional DHB matrix, we identified 148 and 131 endogenous monoisotopic peaks in brain and liver tissue, respectively. In contrast, the ISM DHB+ANI increased the number of annotated ions to 167 in the brain and 145 in liver samples (Figure 1e, 1f). The CHCA matrix performed less effectively, yielding only 86 and 78 annotated monoisotopic peaks in the brain and liver, respectively. However, when used as a solid IM, CHCA+DMA and CHCA+DEA dramatically improved metabolite coverage, reaching 163 and 326 annotated ions in the brain, and 304 and 260 in liver tissue, respectively (Figure 1e, 1f). These results highlight the significant analytical advantage of combining ISMs with LTE, particularly for low molecular weight metabolite detection, within the *m/z* 50–600 Da range.

Given the substantial improvement in metabolite coverage achieved with the ISM CHCA+DEA, we focused our next analyses on brain tissue to validate these findings through high-confidence metabolite identification. Following annotation against the Human Metabolome Database (HMDB) with a 10 ppm mass tolerance, we aimed for level 1 identification confidence by comparing both MS1 adducts and MS/MS fragmentation patterns with authentic standards.

For example, kynurenic acid (KYNA) and taurine were both detected as double potassium adducts, [M+2K–H]^+^, in the standard and brain tissue spectra (Figure 2a, 2b). To rule out misidentification due to isomerism between KYNA and CHCA (both C₁₀H₇NO₃), we verified that the ion at *m/z* 265.9626—assigned to [M+2K–H]^+^ of KYNA—displayed a distinct spatial distribution within the cerebellum, differing from CHCA-related ions (Supplementary Fig. S4a). Furthermore, this ion was absent in matrix-only regions, confirming its tissue-specific origin (Supplementary Fig. S4b).

**Figure 2:**
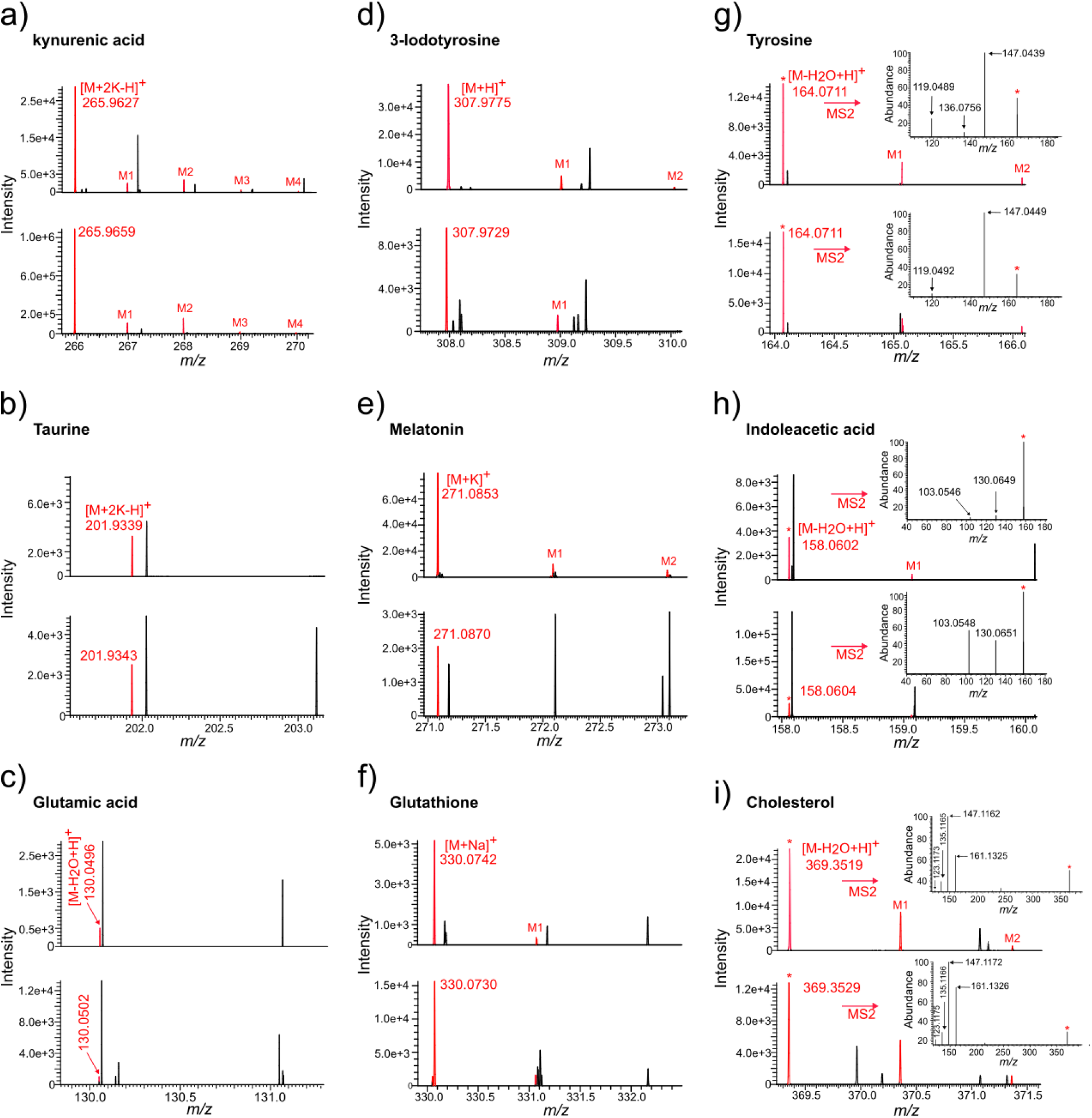
MS spectra of the selected LMWCs: (a) kynurenic acid, (b) taurine, (c) glutamic acid, (d) 3-iodotyrosine, (e) melatonin, and (f) glutathione. MS AND MS/MS spectra of the selected LMWCs: (g) tyrosine, (h) indoleacetic acid, and (i) cholesterol. Spectra on top position were acquired from MLADI-MS analysis of standard compounds and those presented on bottom were obtained from MALDI-MSI of the mouse brain section, using the ISM CHCA+DEA. Peaks related to the selected compound, including isotopes, are colored in red. *: precursor ion for MS/MS analysis.

Further analysis revealed consistent adduct formation between standard and tissue samples for multiple metabolites. For instance, the dehydrated ion [M–H₂O+H]^+^ of glutamic acid, the protonated ion [M+H]^+^ of 3-iodotyrosine (3IAA), the potassium adduct [M+K]^+^ of melatonin, and the sodium adduct [M+Na]^+^ of glutathione all showed reproducible signals in MS1 spectra (Figure 2c–f). Similar results were obtained for additional compounds—including 4-hydroxybenzoic acid (4-HB), alanine, creatine, glutamine, indoleacetic acid, and tyrosine—for which one or more adducts were confirmed across standard and tissue samples (Supplementary Fig. S5–S13). In cases where signal-to-noise ratios allowed, isotopic patterns further corroborated these identifications.

Moreover, high-quality MALDI-MS/MS spectra obtained directly from brain sections enabled structural confirmation of several metabolites. For tyrosine, 3IAA, and cholesterol, the fragmentation patterns closely matched those of the standards (Figure 2g–i). Additionally, MS/MS analysis allowed us to annotate fragments from endogenous compounds such as creatine, dopamine, spermine, N-methylserotonin, choline, and L-carnitine by comparison with reference spectra from HMDB entries, providing deeper insight into their fragmentation patterns (Supplementary Fig. S14).

Combining MS1 adduct verification, isotopic pattern fidelity, and MS2 spectral matching with authentic standards, we confidently identified 73 metabolites below *m/z* 500 from brain tissue with the ISM CHCA+DEA (Supplementary Table S2). This included key neurotransmitters such as acetylcholine, dopamine, L-DOPA, and gamma-aminobutyric acid (GABA). Notably, only 14 of these 73 metabolites were also detected using conventional CHCA or DHB matrices, highlighting the enhanced ionization efficiency afforded by the ISM combined with LTE deposition.

In addition to enhanced molecular coverage, this method also provided significant improvements in spatial fidelity. In MALDI-MSI, particularly for small molecules and metabolites, spatial resolution is often compromised by non-uniform matrix application or the formation of large crystals—issues commonly associated with wet deposition methods. These factors can lead to analyte delocalization due to diffusion from their original locations. In contrast, LTE-deposited solid IMs produce uniform, sub-micrometer-sized crystals and allow precise control of matrix thickness, resulting in a high-purity, homogeneous matrix layer. As shown in Figure 3, MALDI-MSI of 15 selected metabolites at 20 µm resolution yielded sharp, well-defined ion images, demonstrating that the combination of ISMs and LTE enables high-resolution spatial metabolomic mapping of small molecules.

**Figure 3:**
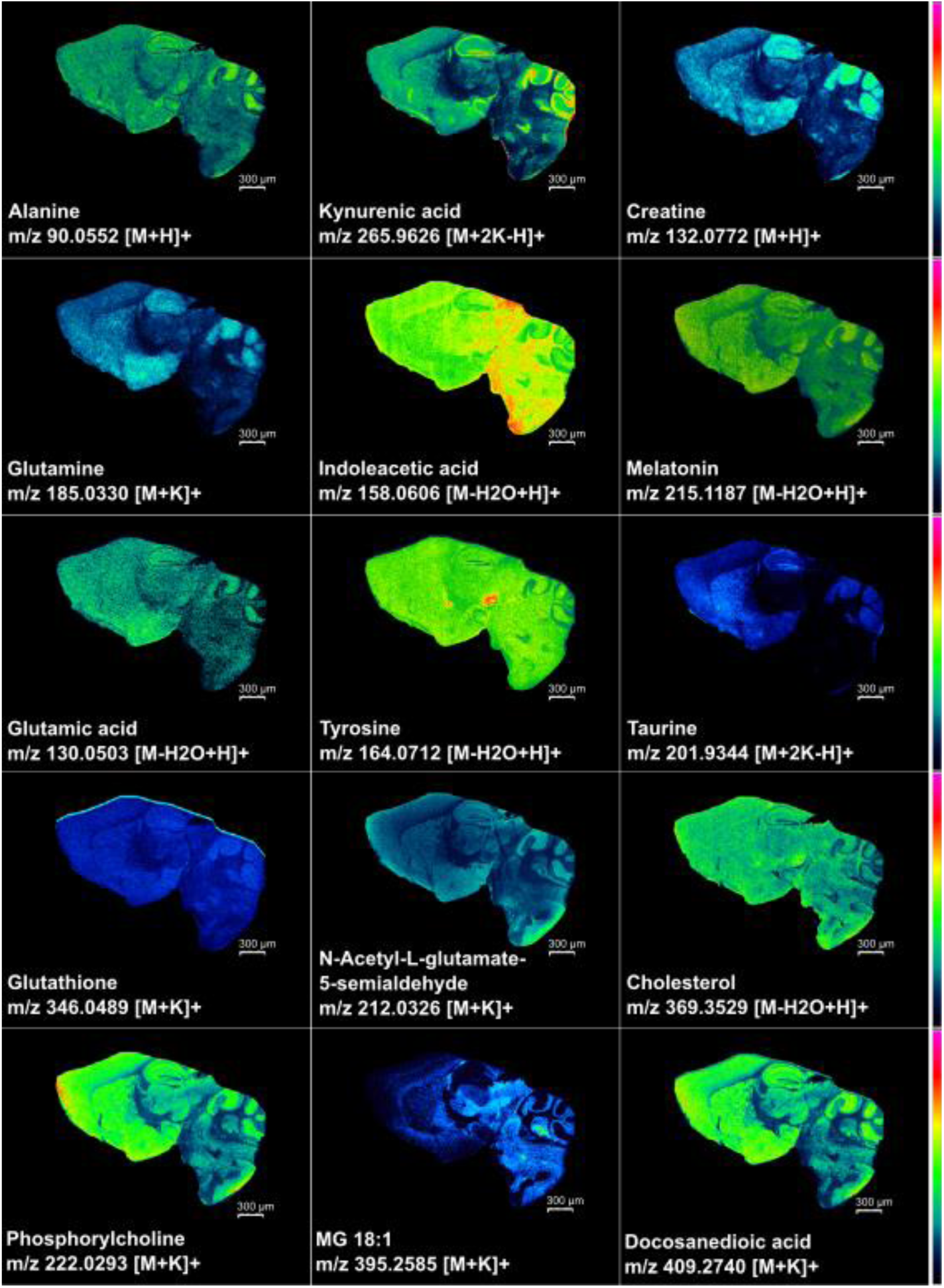
MALDI-MSI of 15 selected metabolites in a mouse sagittal brain section. Ion images were acquired at 20 µm spatial resolution using the ISM CHCA+DEA.

### 3.2. MALDI-MSI of metabolites in mouse pancreatic tissue

To further demonstrate the high-resolution and sensitive capabilities of MALDI-MSI for metabolite imaging, we applied the ISM CHCA+DEA to analyze the metabolic composition of the islets of Langerhans in the pancreas. Each islet consists of a central core of beta cells— accounting for up to 60% of islet mass—surrounded by a mantle of alpha cells (approximately 30%), along with smaller populations of delta, epsilon, and pancreatic polypeptide cells. Alpha cells secrete glucagon, the primary catabolic hormone that increases blood glucose and fatty acid levels, while beta cells produce insulin, which lowers blood glucose by promoting uptake into the liver, muscle, and adipose tissue.

Although MALDI-MSI has been widely used to image pancreatic peptide hormones and proteins [33], its application to lipids and small metabolites within islets has remained limited. Prior studies localized only a few lipid species—such as specific lysophosphatidylcholines (LPCs), phosphatidylcholines (PCs), and GM3 gangliosides—within murine islet cells [34], findings that were also supported by nano-DESI MSI [35]. For small metabolites, compounds like glucose 6-phosphate, fatty acids, ADP, cholesterol sulfate, and 3-O-sulfogalactosylceramide have shown considerable intra- and inter-islet variability [36].

Here, in addition to the previously described LPC and PC species, we show ion images acquired at 25 µm spatial resolution (Figure 4) revealing the distinct localization of small metabolites to the islets of Langerhans. These include ions at *m/z* 215.1185 and *m/z* 297.1266, identified as melatonin and phenylalanyl-methionine, respectively, were enriched along the periphery of the islets. Conversely, metabolites such as 3-methylglutarylcarnitine (*m/z* 213.1413), 3-iodotyrosine (*m/z* 307.9779), and 6-beta-hydroxycortisol (*m/z* 417.1726) were more uniformly distributed throughout the islets, with no significant difference between core and edge regions.

**Figure 4:**
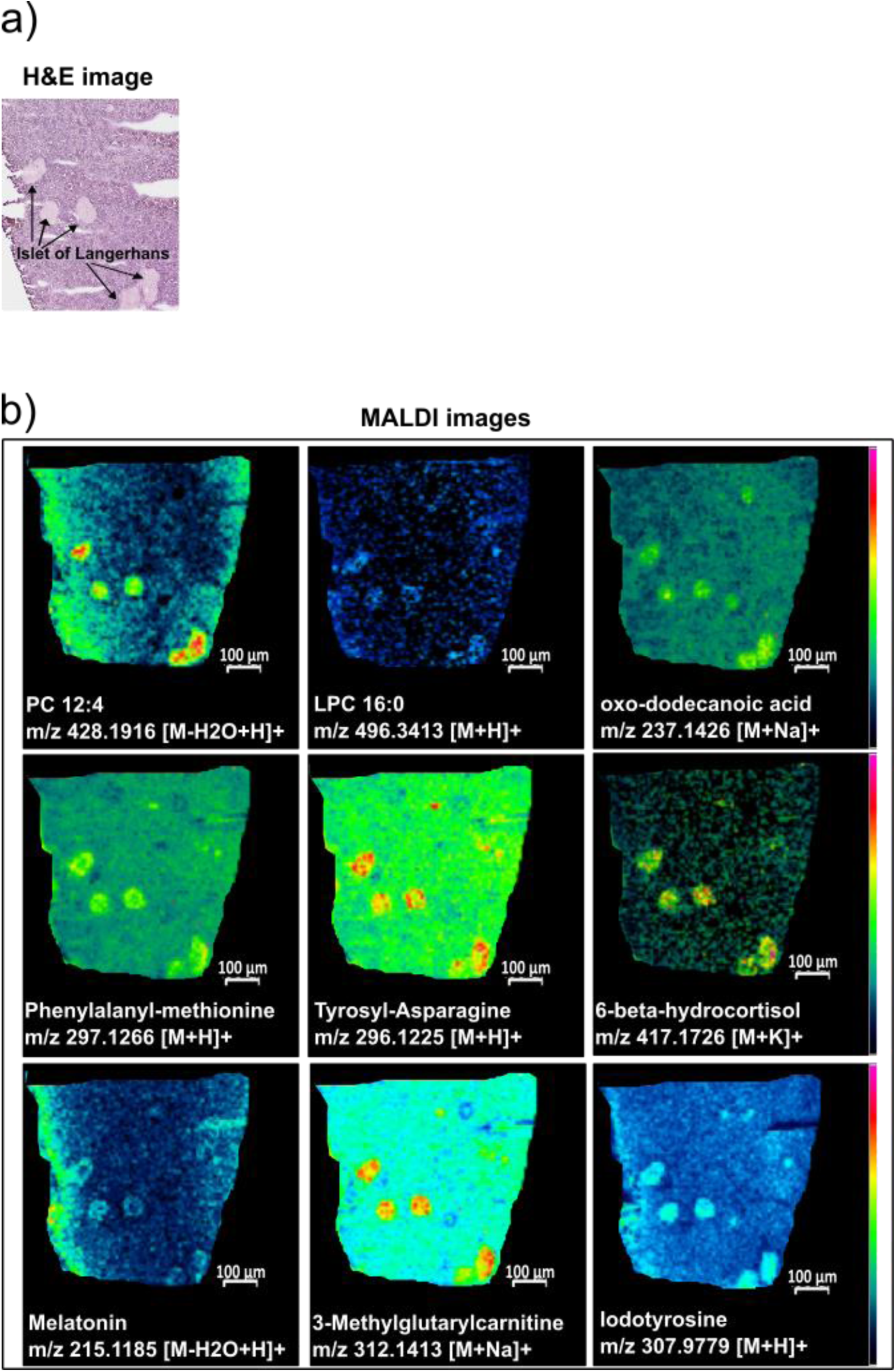
Spatial localization of small metabolites in mouse pancreas. (a) H&E-stained image of pancreas tissue with five islets of Langerhans indicated by black arrows. (b) MALDI-MSI ion images of nine selected metabolites acquired at 25 µm spatial resolution using the ISM CHCA+DEA, showing specific localization to the islets of Langerhans.

Melatonin plays a key regulatory role in insulin and glucagon secretion through its interaction with MTNR1A and MTNR1B receptors, which are expressed in both α- and β-cells [37]. Although MTNR1A protein expression has previously been localized to peripherally positioned β-cells, to our knowledge, this is the first report of direct detection of melatonin within pancreatic islets. This finding provides MALDI-MSI as a useful and sensitive technique to potentially link circadian rhythms and metabolic homeostasis, highlighting melatonin’s dual role in modulating both insulin and glucagon output.

Iodotyrosine is a precursor in TH synthesis, and TH receptors (TRα/TRβ) are expressed in pancreatic islets [38,39]. The presence of iodotyrosine implies local TH metabolism, which could fine-tune insulin release in response to nutrient availability.

### Conclusions

We present a novel strategy for enhancing MALDI-MSI of low molecular weight compounds (LMWCs): the application of solid ionic matrices (solid-IMs) via low-temperature thermal evaporation (LTE). Compared to conventional matrices such as CHCA and DHB, solid-IMs offer clear advantages: (1) a significant reduction in matrix-related ion signals, minimizing spectral interference with analyte peaks, and (2) improved sensitivity and signal intensities for tissue-derived ions.

Using solid IM CHCA+DEA, we demonstrated reduced matrix cluster formation and enhanced ionization efficiency in mouse brain tissue, enabling the detection and annotation of 73 small metabolites (m/z < 500). The resulting MALDI-MSI data at 20 µm resolution provided high-quality images with clear spatial localization of metabolites within brain regions.

Finally, while MALDI-MSI has traditionally focused on imaging peptide hormones [40–42] and lipids [34,35] in the pancreas, application of CHCA+DEA with LTE enabled the detection and specific localization of metabolites—such as melatonin and 3-iodotyrosine—within the islets of Langerhans. These findings offer valuable insights into the pancreatic metabolic landscape and open new opportunities for advancing our understanding of islet biology, function, and disease.

## Acknowledgements

TM acknowledges the financial support of the Universitat Rovira i Virgili through the predoctoral grant ref. PRE2019-089374. OY acknowledges the financial support of grant PID2022-136226OB-I00 funded by MICIU/AEI/10.13039/501100011033 and by ERDF/EU, and grant TED2021-132635B-I00 funded by MICIU/AEI/10.13039/501100011033 and by the European Union NextGenerationEU/PRTR.

