## Supplementary information for "Solid Ionic Matrices applied via Low-Temperature Evaporation enable High-Resolution and Sensitive MALDI Imaging of Metabolites"

**Table S1:** Evaluation of ionic matrices, in terms of structure, physical state, and the optimised temperature for low-temperature thermal evaporation deposition

| Matrix | Structure | State of aggregation (at 25ºC) | LTE temperature |
|---|---|---|---|
| CHCA | 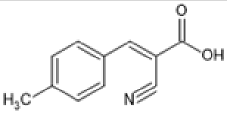 | Solid | 100 ºC |
| CHCA+ANI | 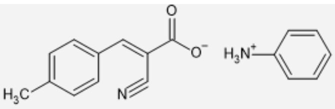 | Solid | 112 ºC |
| CHCA+DEA | 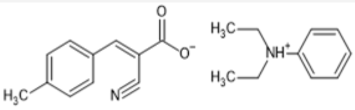 | Solid | 115 ºC |
| CHCA+DMA | 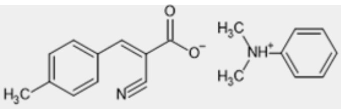 | Solid | 115 ºC |
| DHB | 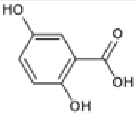 | Solid | 80 ºC |
| DHB+ANI | 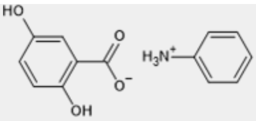 | Solid | 90 ºC |
| DHB+DEA | 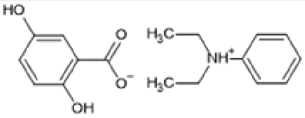 | Liquid | / |
| DHB+DMA | 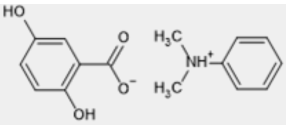 | Liquid | / |

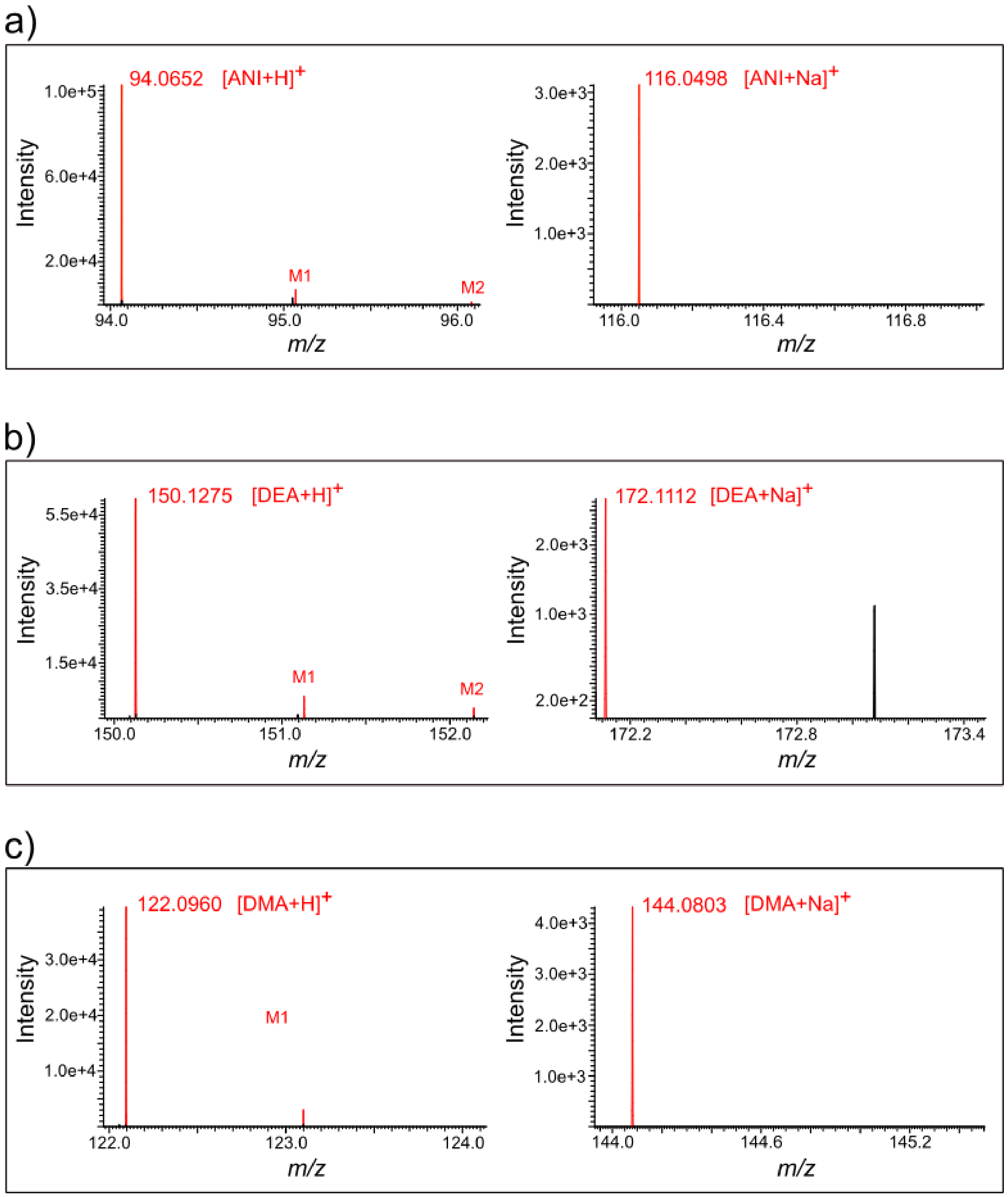


**Figure S1:** MALDI-MS spectra acquired from matrix only section showing the detection of peaks from the base components of the solid ionic matrix; (a) the protonated ion (*m/z* 94.0652) and the sodium adduct ion (*m/z* 116.0498) of aniline added to DHB matrix; (b) the protonated ion (*m/z* 150.1275) and the sodium adduct ion (*m/z* 172.1112) of N,N-diethylaniline added to CHCA matrix; (c) the protonated ion (*m/z* 122.0960) and the sodium adduct ion (*m/z* 144.0803) of N,N-dimethylaniline added to CHCA matrix.

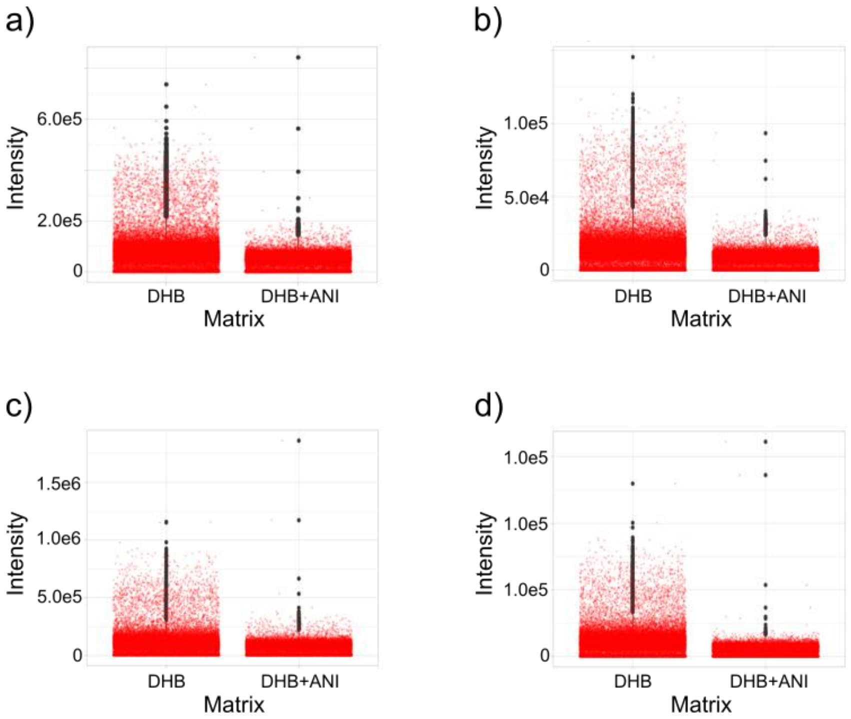


**Figure S 2:** Plot intensity of the key DHB related matrix ions detected using DHB matrix and the solid ionic matrix DHB+ANI; (a) *m/z* 137.0239 ($[M\text{-}H_2O+H]^+$), (b) *m/z* 155.0345 ($[M+H]^+$), (c) *m/z* 273.0399 ($[2M\text{-}2(H_2O)+H]^+$), and (d) *m/z* 409.0559 ($[3M\text{-}3(H_2O)+H]^+$).

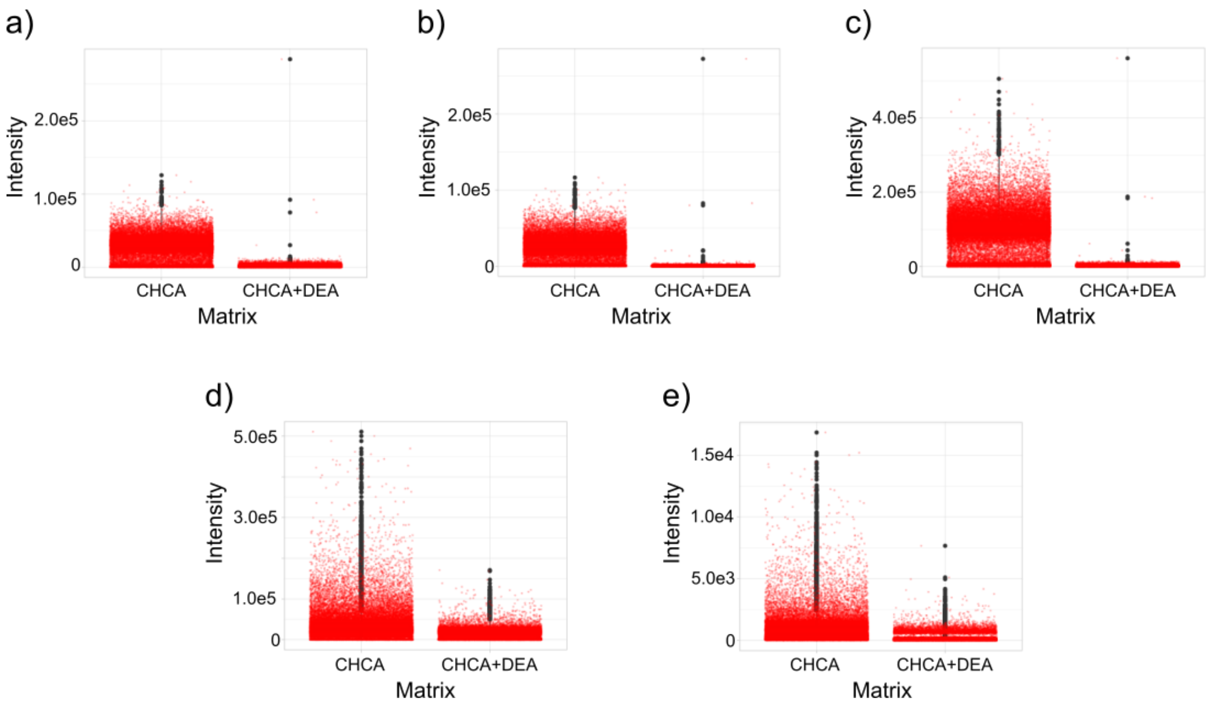


**Figure S 3:** Plot intensity of the key CHCA related matrix ions detected using CHCA matrix and the solid ionic matrix CHCA+DEA; (a) *m/z* 146.0613 ($[M\text{-}CO+H]^+$), (b) *m/z* 172.0394 ($[M\text{-}H2O+H]^+$), (c) *m/z* 190.0511 ($[M+H]^+$), (d) *m/z* 212.0340 ($[M+Na]^+$), and (e) *m/z* 228.0067 ($[M+K]^+$).

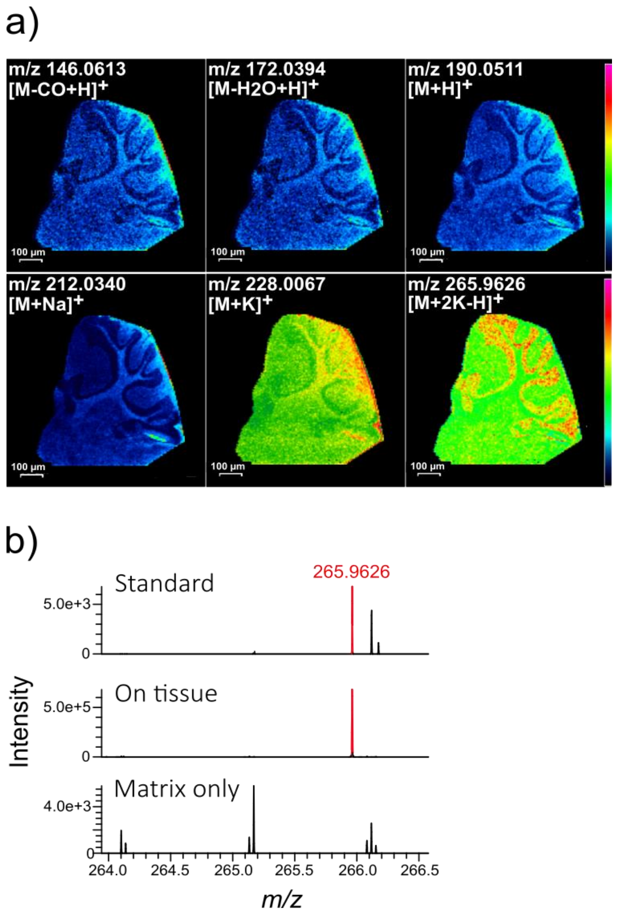


**Figure S 4:** (a) MALDI images of mouse cerebellum acquired at 30 µm, using the solid ionic matrix CHCA+DEA, of the detected ions at *m/z* 146.0613, *m/z* 172.0394, *m/z* 190.0511, *m/z* 212.0340, *m/z* 228.0067, and *m/z* 265.9626 annotated as $[M\text{-}CO+H]^+$, $[M\text{-}H_2O+H]^+$, $[M+H]^+$, $[M+Na]^+$, $[M+K]^+$, and $[M+2K\text{-}H]^+$ respectively. (b) MLADI-MS spectra showing the detection the of the peak at *m/z* 265.9626 from the standard solution and from the brain tissue, but not from the matrix only measurements.

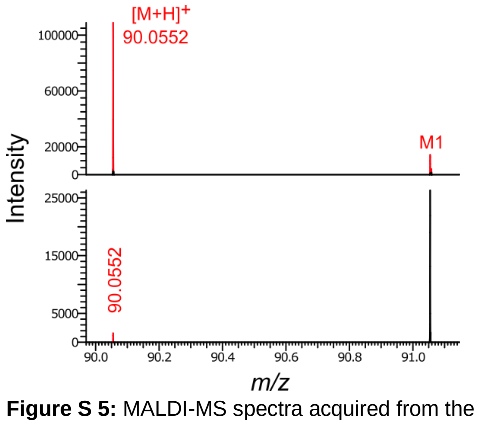


**Figure S 5:** MALDI-MS spectra acquired from the dried-droplet of alanine standard analysis (top position) and brain section imaging (bottom position) using the ISM CHCA+DEA.

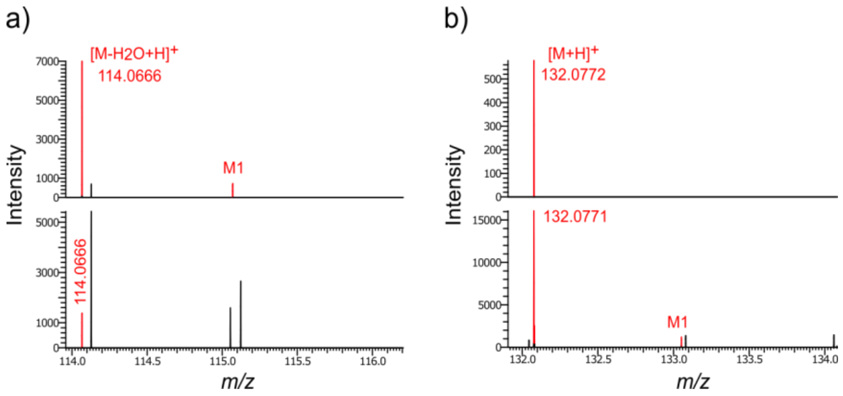


**Figure S 6:** MALDI-MS spectra acquired from the dried-droplet of creatine standard analysis (top position) and brain section imaging (bottom position) using the ISM CHCA+DEA: (a) zoomed in at the peak *m/z* 114.06 ([M-$H_2$O+H]$^+$) and (b) zoomed in at the peak *m/z* 132.07 ([M+H]$^+$).

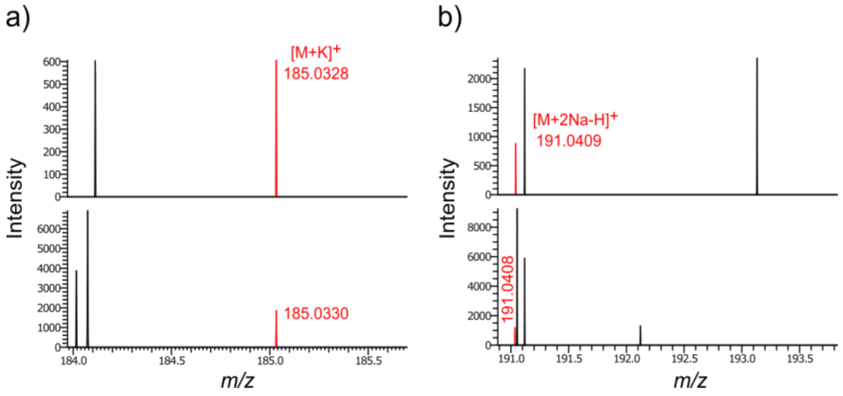


**Figure S 7:** MALDI-MS spectra acquired from the dried-droplet of glutamine standard analysis (top position) and brain section imaging (bottom position) using the ISM CHCA+DEA: (a) zoomed in at the peak *m/z* 185.03 ([M+K]$^+$) and (b) zoomed in at the peak *m/z* 191.04 ([M+2Na-H]$^+$).

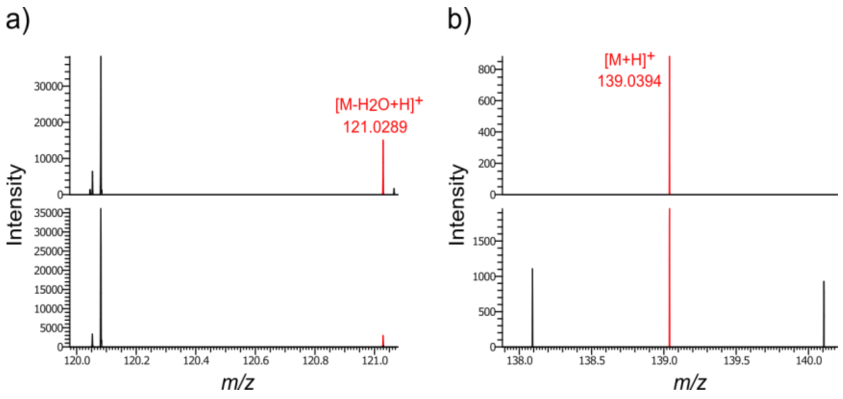


**Figure S 8:** MALDI-MS spectra acquired from the dried-droplet of 4-hydroxybenzoic acid standard analysis (top position) and brain section imaging (bottom position) using the ISM CHCA+DEA: (a) zoomed in at the peak *m/z* 121.0289 ([M-$H_2$O+H]$^+$) and (b) zoomed in at the peak *m/z* 139.0394 ([M+H]$^+$).

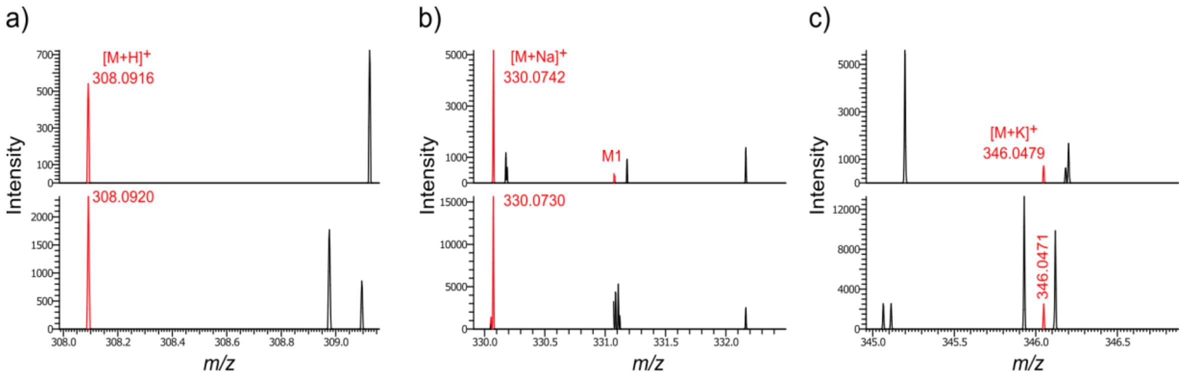


**Figure S 9:** MALDI-MS spectra acquired from the dried-droplet of glutathione standard analysis (top position) and brain section imaging (bottom position) using the ISM CHCA+DEA: (a) zoomed in at the peak *m/z* 308.09 ($[M+H]^+$); (b) zoomed in at the peak *m/z* 330.0742 ($[M+Na]^+$); and (c) zoomed in at the peak *m/z* 346.05 ($[M+K]^+$).

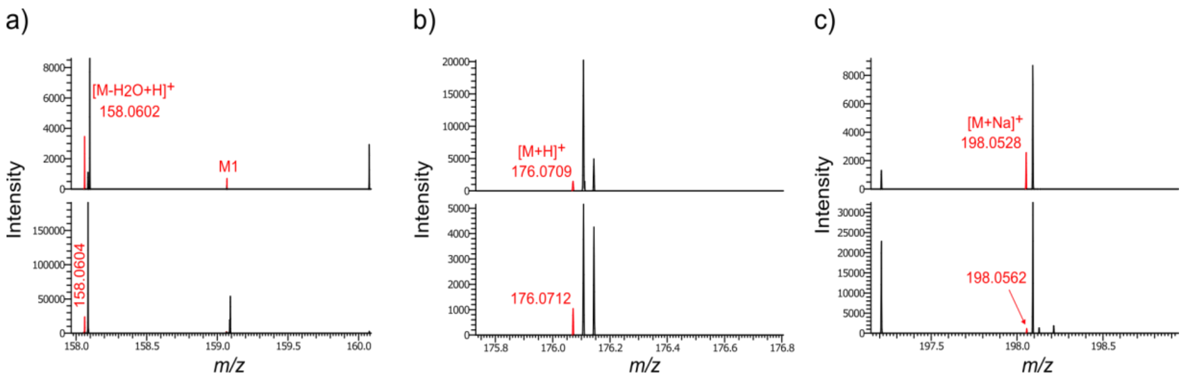


**Figure S 10:** MALDI-MS spectra acquired from the dried-droplet of indoleacetic acid standard analysis (top position) and brain section imaging (bottom position) using the ISM CHCA+DEA: (a) zoomed in at the peak *m/z* 158.06 ($[M-H_2O+H]^+$); (b) zoomed in at the peak *m/z* 176.07 ($[M+H]^+$); and (c) zoomed in at the peak *m/z* 198.05 ($[M+Na]^+$).

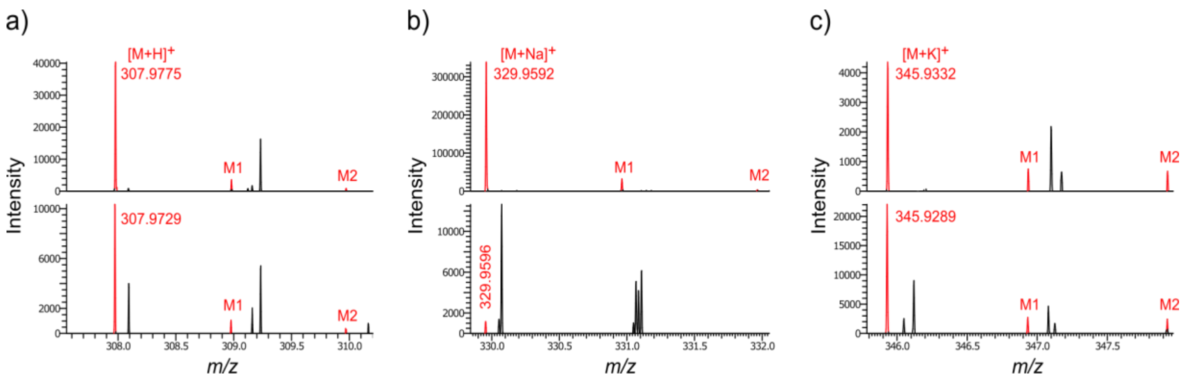


**Figure S 11:** MALDI-MS spectra acquired from the dried-droplet of 3-iodotyrosine standard analysis (top position) and brain section imaging (bottom position) using the ISM CHCA+DEA: (a) zoomed in at the peak *m/z* 307.09 ($[M+H]^+$); (b) zoomed in at the peak *m/z* 329.96 ($[M+Na]^+$); and (c) zoomed in at the peak *m/z* 345.93 ($[M+K]^+$).

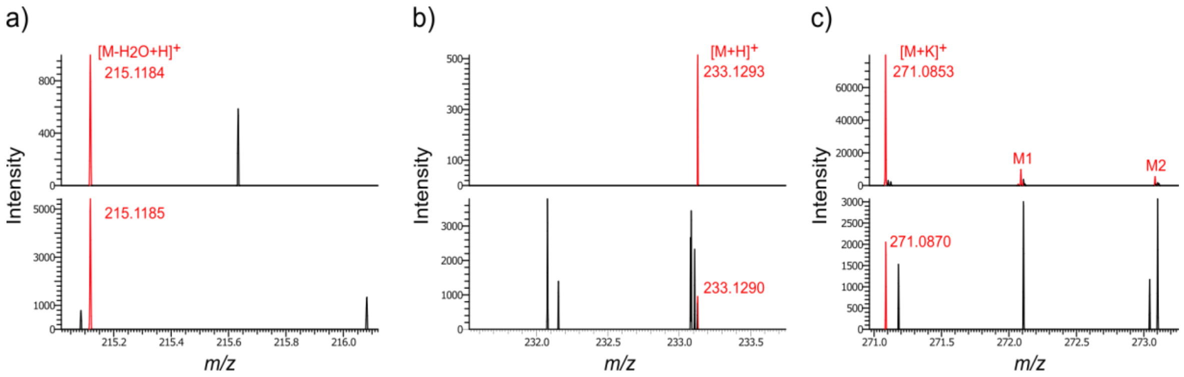


**Figure S 12:** MALDI-MS spectra acquired from the dried-droplet of melatonin standard analysis (top position) and brain section imaging (bottom position) using the ISM CHCA+DEA: (a) zoomed in at the peak *m/z* 215.12 ($[M-H_2O+H]^+$); (b) zoomed in at the peak *m/z* 233.13 ($[M+Na]^+$); and (c) zoomed in at the peak *m/z* 271.08 ($[M+K]^+$).

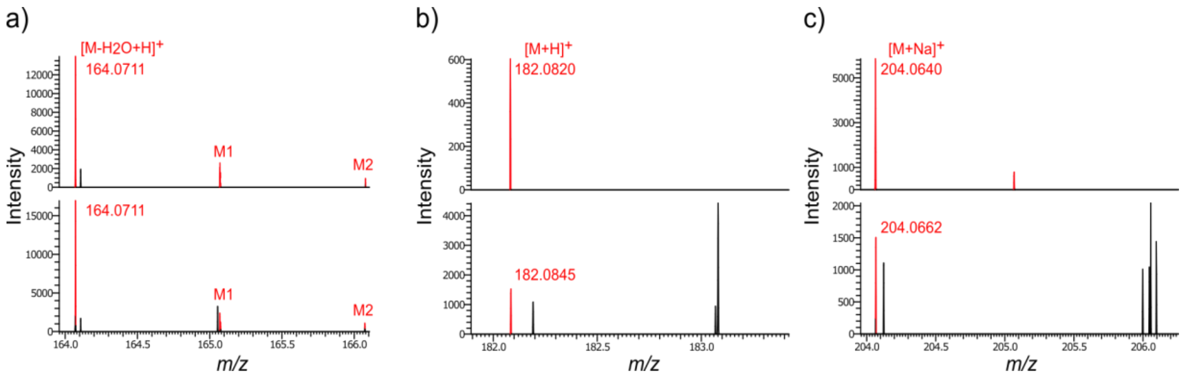


**Figure S 13:** MALDI-MS spectra acquired from the dried-droplet of tyrosine standard analysis (top position) and brain section imaging (bottom position) using the ISM CHCA+DEA: (a) zoomed in at the peak *m/z* 164.07 ($[M-H_2O+H]^+$); (b) zoomed in at the peak *m/z* 182.08 ($[M+H]^+$); and (c) zoomed in at the peak *m/z* 204.06 ($[M+Na]^+$).

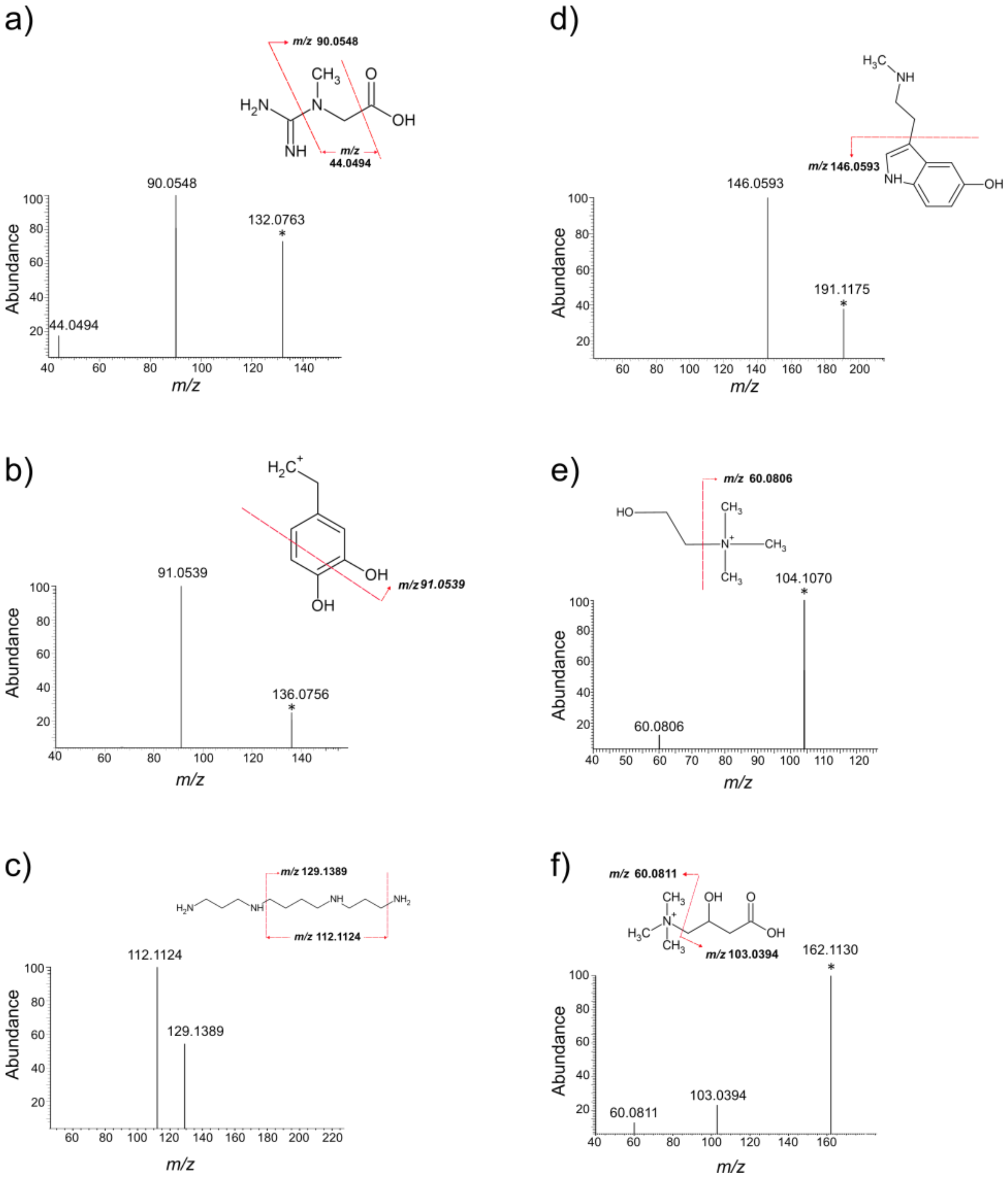


**Figure S14:** MALDI-MS/MS spectra acquired from the brain section imaging using the ISM CHCA+DEA, with the proposed fragmentation pathways of: (a) creatine; (b) dopamine; (c) spermine; (d) n-methylserotonin; (e) choline; and (f) carnitine.

**Table S 2:** List of LMWCs (*m/z* < 500 Da) annotated from MALDI-MSI of mouse brain using the ISM CHCA+DEA.

| Compound | *m/z* | ppm error (+/-) | Adduct type |
|---|---|---|---|
| Acetaminophen | 134.0605 | 0.04 | $[M-H_2O+H]^+$ |
| Acetic acid | 136.9403 | 1.25 | $[M+2K-H]^+$ |
| N-Acetyl-L-aspartic acid | 251.968 | 1.76 | $[M+2K-H]^+$ |
| L-Acetylcarnitine * | 204.1231 | 0.52 | $[M+H]^+$ |
| Acetylcholine | 146.1175 | 4.05 | $[M+H]^+$ |
| Acetyl-N-formyl-5-methoxykynurenamine | 303.0774 | 9.15 | $[M+H]^+$ |
| N-Acetyl-L-glutamate 5-semialdehyde | 212.0326 | 2.9 | $[M+K]^+$ |
| Adenosine monophosphate | 330.0575 | 8.32 | $[M-H_2O+H]^+$ |
| Alanine | 90.0552 | 0.0002 | $[M+H]^+$ |
| L-Alloisoleucine | 114.0917 | 1.55 | $[M-H_2O+H]^+$ |
| Arginine * | 175.1196 | 0.6 | $[M+H]^+$ |
| L-Aspartic acid | 134.0452 | 0.44 | $[M+H]^+$ |
| L-carnitine * | 162.1130 | 3.0 | $[M+H]^+$ |
| Cholesterol * | 369.3529 | 2.31 | $[M-H_2O+H]^+$ |
| Choline * | 104.1071 | 1.99 | $[M+H]^+$ |
| Citrulline | 198.0866 | 6.17 | $[M+Na]^+$ |
| Creatine * | 132.0772 | 0.38 | $[M+H]^+$ |
| | 114.0665 | 1.45 | $[M-H_2O+H]^+$ |
| L-Cystathionine | 223.0762 | 4.54 | $[M+H]^+$ |
| 7-Dehydrocholesterol * | 367.3371 | 1.9 | $[M-H_2O+H]^+$ |
| Diacetyl * | 162.9559 | 0.89 | $[M+2K-H]^+$ |

| | | | |
|---|---|---|---|
| Dihydroxyphenyl-Acetaldehyde (DOPAL) | 135.0445 | 0.71 | $[M-H_2O+H]^+$ |
| Dihydroxyphenylglycol (DOPEG) | 105.0702 | 1.7 | $[M-H_2O+H]^+$ |
| Docosahexaenoic acid | 367.2045 | 1.74 | [M+K]+ |
| Docosanedioic acid | 409.2740 | 3.3 | $[M+Na]^+$ |
| DOPA | 180.0662 | 1.1 | $[M-H_2O+H]^+$ |
| Dopamine | 136.0762 | 0.18 | $[M-H_2O+H]^+$ |
| Erythronic acid | 137.0462 | 9.07 | $[M+H]^+$ |
| gamma-Aminobutyric acid (GABA) | 104.0709 | 1.99 | $[M+H]^+$ |
| | 179.9826 | 1.34 | $[M+2K-H]^+$ |
| Ethanolamine | 62.0602 | 4.67 | $[M+H]^+$ |
| Glutamic acid * | 130.0503 | 0.64 | $[M-H_2O+H]^+$ |
| Glutamine | 185.033 | 0.81 | $[M+K]^+$ |
| | 191.0409 | 1.36 | $[M+2Na-H]^+$ |
| Glutathione | 308.0929 | 4.33 | $[M+H]^+$ |
| | 330.0742 | 4.66 | $[M+Na]^+$ |
| | 346.0489 | 4.05 | $[M+K]^+$ |
| Hippuric acid | 162.0555 | 0.54 | $[M-H_2O+H]^+$ |
| 4-Hydroxybenzoic acid (4-HB) | 121.0288 | 0.92 | $[M-H_2O+H]^+$ |
| | 139.0395 | 0.09 | $[M+H]^+$ |
| Hydroxycholesterol * | 385.3478 | 2.01 | $[M-H_2O+H]^+$ |
| 5-Hydroxyindoleacetic acid | 192.0662 | 0.94 | $[M+H]^+$ |
| | 174.0555 | 0.01 | $[M-H_2O+H]^+$ |

| | | | |
|---|---|---|---|
| 6-Hydroxymelatonin | 231.1135 | 0.98 | $[M-H_2O+H]^+$ |
| | 287.0825 | 9.63 | $[M+K]^+$ |
| Hydroxypropionylcarnitine | 272.0925 | 9.34 | $[M+K]^+$ |
| 5-Hydroxytryptophol | 160.0763 | 0.41 | $[M-H_2O+H]^+$ |
| Indole | 118.0464 | 0.01 | $[M+H]^+$ |
| Indoleacetic acid | 158.0606 | 0.61 | $[M-H_2O+H]^+$ |
| | 176.0712 | 0.51 | $[M+H]^+$ |
| | 198.0528 | 1.42 | $[M+Na]^+$ |
| Iodotyrosine | 307.9775 | 0.95 | $[M+H]^+$ |
| | 329.9592 | 1.78 | $[M+Na]^+$ |
| | 345.9332 | 0.95 | $[M+K]^+$ |
| 7-Ketocholesterol | 401.3428 | 2.34 | $[M+H]^+$ |
| | 383.3313 | 0.09 | $[M-H_2O+H]^+$ |
| Kynurenic acid | 265.9626 | 1.69 | $[M+2K-H]^+$ |
| Kyotorphin | 376.1394 | 2.08 | $[M+K]^+$ |
| Lactic acid | 166.9506 | 0.09 | $[M+2K-H]^+$ |
| Melatonin | 215.1184 | 1.05 | $[M-H_2O+H]^+$ |
| | 233.1292 | 1.05 | $[M+H]^+$ |
| | 271.0853 | 1.15 | $[M+K]^+$ |
| 5-Methoxytryptophol | 174.0919 | 0.5 | $[M-H_2O+H]^+$ |
| 3-Methoxytyramine | 150.0919 | 0.22 | $[M-H_2O+H]^+$ |
| Methylguanidine | 74.0715 | 4.17 | $[M+H]^+$ |

| | | | |
|---|---|---|---|
| Methylimidazoleacetic acid | 123.0557 | 1.01 | $[M-H_2O+H]^+$ |
| N(6)-Methyllysine | 143.1184 | 0.05 | $[M-H_2O+H]^+$ |
| (S)-N-Methylsalsolinol | 232.0761 | 9.21 | $[M+K]^+$ |
| | 176.1076 | 0.43 | $[M-H_2O+H]^+$ |
| N-Methylserotonin | 191.1175 | 0.95 | $[M+H]^+$ |
| MG 18:1 | 395.2585 | 2.7 | $[M+K]^+$ |
| Neuraminic acid | 290.0823 | 9.67 | $[M+Na]^+$ |
| O-Phosphoethanolamine | 124.0162 | 0.85 | $[M-H_2O+H]^+$ |
| | 217.9385 | 1.98 | $[M+2K-H]^+$ |
| Palmitoylethanolamide * | 282.2802 | 1.94 | $[M-H_2O+H]^+$ |
| Parathion | 292.0379 | 9.98 | $[M+H]^+$ |
| Pentosidine | 417.1611 | 9.8 | $[M+K]^+$ |
| Phenylacetic acid | 137.0601 | 1.07 | $[M+H]^+$ |
| | 119.0495 | 1.08 | $[M-H_2O+H]^+$ |
| Phosphodimethylethanolamine | 245.9699 | 1.85 | $[M+2K-H]^+$ |
| Phosphorylcholine * | 184.0733 | 3 | $[M+H]^+$ |
| Pipecolic acid | 205.9986 | 0.38 | $[M+2K-H]^+$ |
| Proline | 154.0267 | 2.06 | $[M+K]^+$ |
| Propionylcarnitine * | 240.1221 | 4.16 | $[M+Na]^+$ |
| Propionylcholine | 198.0915 | 4.17 | $[M+K]^+$ |
| Pyridoxal | 150.0554 | 0.32 | $[M-H_2O+H]^+$ |
| Pyroglutamic acid | 112.0397 | 1.35 | $[M-H_2O+H]^+$ |
| Spermine * | 203.2237 | 1.04 | $[M+H]^+$ |
| Taurine | 163.9784 | 0.75 | $[M+K]^+$ |

| | | | |
|---|---|---|---|
| | 201.9344 | 1.16 | $[M+2K-H]^+$ |
| Trimethyl-L-lysine | 171.1498 | 0.7 | $[M-H_2O+H]^+$ |
| Tyrosine | 164.0712 | 0.59 | $[M-H_2O+H]^+$ |
| | 182.0820 | 0.76 | $[M+H]^+$ |
| | 204.0662 | 2.65 | $[M+Na]^+$ |

* compounds detected using the conventional matrix (CHCA or DHB).

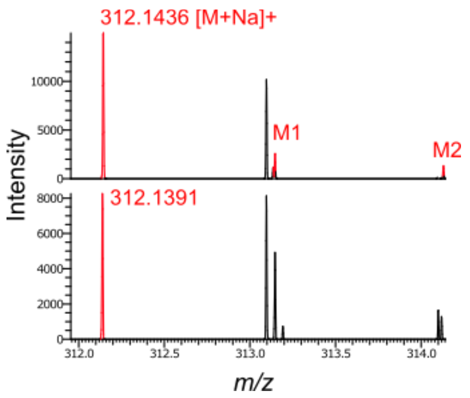


**Figure S 15:** MALDI-MS spectra acquired from the dried-droplet of 3-methyl-glutarylcarnitine standard analysis (top position) and mouse pancreatic section imaging(bottom position) using the ISM CHCA+DEA.

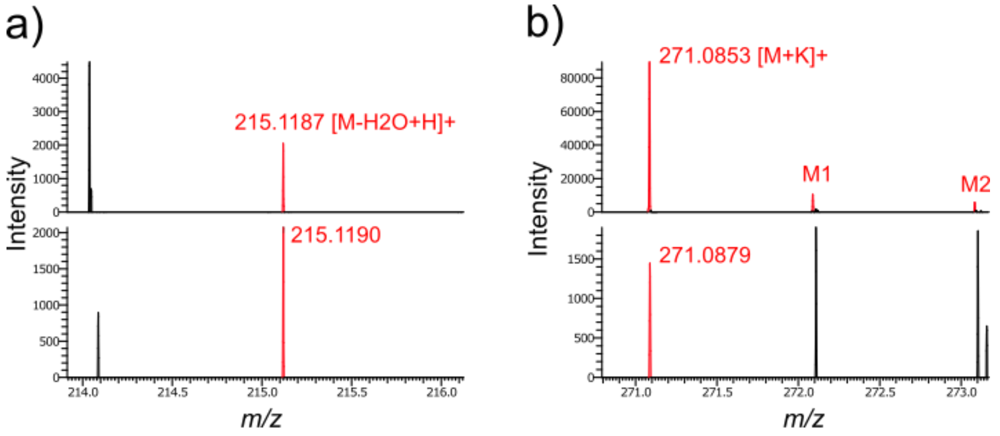


**Figure S 16:** MALDI-MS spectra acquired from the dried-droplet of melatonin standard analysis (top position) and mouse pancreatic section imaging (bottom position) using the ISM CHCA+DEA: (a) zoomed in at the peak *m/z* 215.12 ($[M-H_2O+H]^+$) and (b) zoomed in at the peak *m/z m/z* 271.08 ($[M+K]^+$).

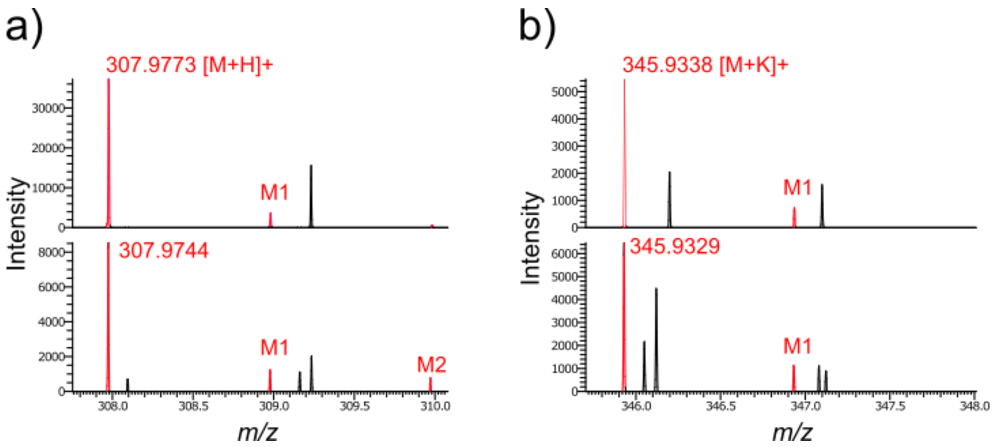


**Figure S 17:** MALDI-MS spectra acquired from the dried-droplet of 3-iodotyrosine standard analysis (top position) and mouse pancreatic section imaging (bottom position) using the ISM CHCA+DEA: (a) zoomed in at the peak *m/z* 307.09 ($[M+H]^+$) and (b) zoomed in at the peak *m/z* 345.93 ($[M+K]^+$).

.
